# The conservation status and population decline of the African penguin deconstructed in space and time

**DOI:** 10.1101/2020.01.15.907485

**Authors:** Richard B. Sherley, Robert J. M. Crawford, Andrew D. de Blocq, Bruce M. Dyer, Deon Geldenhuys, Christina Hagen, Jessica Kemper, Azwianewi B. Makhado, Lorien Pichegru, Desmond Tom, Leshia Upfold, Johan Visagie, Lauren J. Waller, Henning Winker

## Abstract

Understanding changes in abundance is crucial for conservation, but population growth rates often vary over space and time. We use 40 years of count data (1979–2019) and Bayesian state-space models to assess the African penguin *Spheniscus demersus* population under IUCN Red List Criterion A. We deconstruct the overall decline in time and space to identify where urgent conservation action is needed. The global African penguin population met the threshold for *Endangered* with a high probability (97%), having declined by almost 65% since 1989. An historical low of ~17,900 pairs bred in 2019. Annual declines were faster in the South African population (−4.2%, highest posterior density interval, HPDI: −7.8 to −0.6%) than the Namibian one (−0.3%, HPDI: −3.3 to +2.6%), and since 1999 were almost 10% at South African colonies north of Cape Town. Over the 40-year period, the Eastern Cape colonies went from holding ~25% of the total penguin population to ~40% as numbers decreased more rapidly elsewhere. These changes coincided with an altered abundance and availability of the main prey of African penguins. Our results underline the dynamic nature of population declines in space as well as time and highlight which penguin colonies require urgent conservation attention.

## 1. Introduction

Seabirds are considered to be the most threatened group of birds in the world (Croxall et al., 2012); globally their populations may have declined by >70% since 1950 (Paleczny, Hammill, Karpouzi, & Pauly, 2015). Seabirds face a number of threats both on land in their colonies, like invasive alien species and disturbance, and in the oceans, such as bycatch and competition with fisheries (Dias et al., 2019). Seven seabird species breed only within the influence of the Benguela upwelling ecosystem of Southern Africa (Angola, Namibia and South Africa). Five of these endemics are listed in a threatened category (Vulnerable or worse) on the International Union for Conservation of Nature (IUCN) Red List, including the African penguin *Spheniscus demersus* which was first listed as Endangered in 2010 (Crawford et al., 2011).

The African penguin breeds, or has bred, at 32 island and mainland colonies between central Namibia (Hollamsbird Island) and South Africa’s Eastern Cape province (Bird Island; Figure 1) (Crawford, Kemper, & Underhill, 2013). The breeding colonies are clustered in three core groups, Namibia, South Africa’s Western Cape, and South Africa’s Eastern Cape, each separated from another by c. 600 km (Figure 1). Although the total population at the turn of the 20^th^ century is not known, there may have been as many as 1.5–3.0 million individuals across the species range and 0.3 million pairs on Dassen Island alone (Frost, Siegfried, & Cooper, 1976; Shannon & Crawford, 1999; Crawford, Underhill, Upfold, & Dyer, 2007). By 1956, only an estimated 0.3 million individuals remained, and the population has more or less declined consistently since then apart from a period in the late-1990s and early-2000s when numbers in the Western Cape briefly recovered (Crawford et al., 2011). This population change since the 1950s has been attributed to a number of top-down and bottom-up processes, including historical egg collecting and guano scraping, changes in the abundance and distribution of their main prey (sardine *Sardinops sagax* and anchovy *Engraulis encrasicolus*), pollution, habitat loss and modification, predation on land in their colonies and at sea, competition with fisheries, and climate change (Frost et al., 1976; Crawford, 2007; Sherley et al., 2017; Crawford, Makhado, & Oosthuizen, 2018).

**Figure 1.**
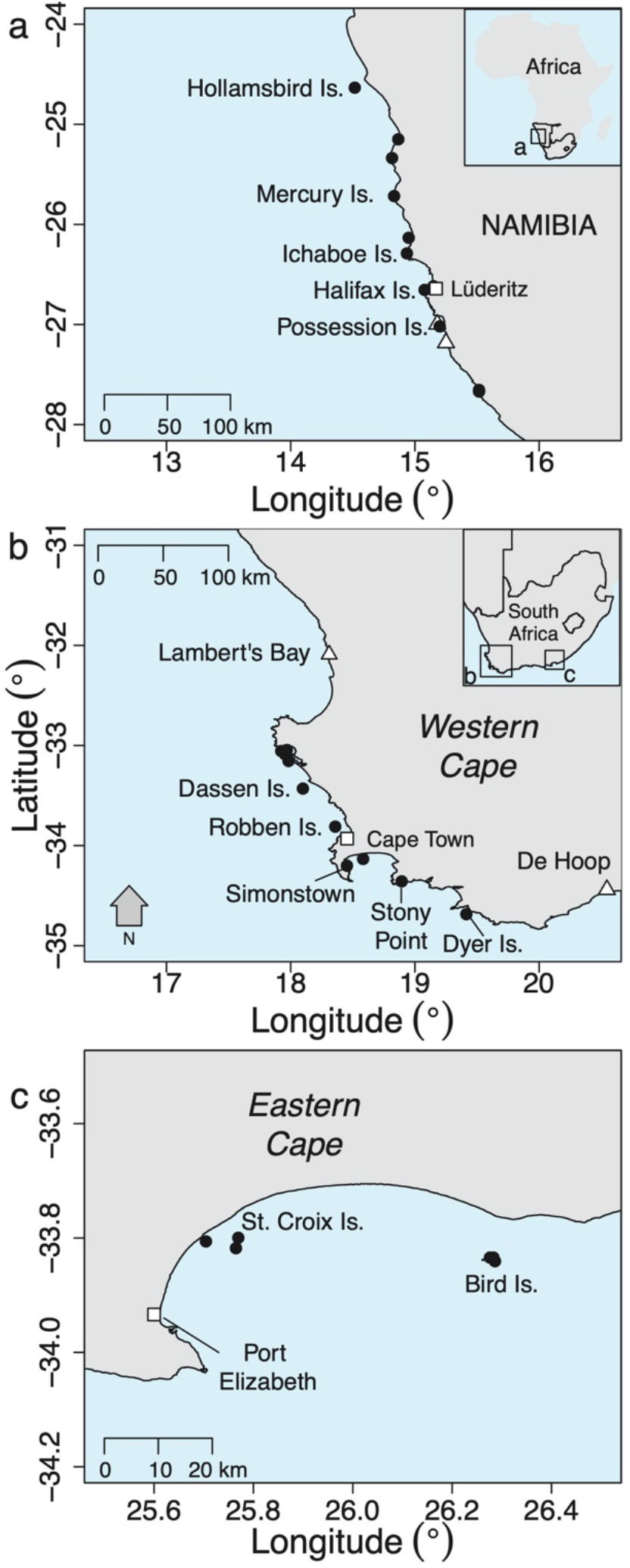
The 28 extant (●) and 4 extinct (Δ) breeding colonies of the African penguin in South Africa and Namibia. Colonies mentioned in the text are named, as are the major towns and cities (□) in each region.

African penguin breeding populations have been counted at all major colonies in South Africa since 1979 and at the four major colonies in Namibia since 1985 (Crawford et al., 2013). Here, we use these count data and a generalized Bayesian state-space tool for estimating extinction risk estimates under IUCN Red List Criterion A (Just Another Red List Assessment [JARA], Sherley et al., 2019b; Winker & Sherley, 2019) to assess the current status of the African penguin population at a global scale. We then deconstruct the overall decline in time and space to identify the regional populations most in need of urgent conservation action. Finally, we review the threats faced by the species and identify interventions needed to secure the species’ conservation in light of our findings.

## 2. Materials and Methods

### 2.1 Penguin count data

In South Africa, the numbers of occupied nest sites of African penguins were counted at most extant breeding colonies sporadically between 1979 and 1991 and annually thereafter (Shelton, Crawford, Cooper, & Brooke, 1984; Crawford et al., 2011). We used counts from 18 localities where penguins bred in South Africa for more than 5 of the 41 years from 1979 to 2019 (Figure S1, Appendix 1). Of a possible 737 annual counts, 472 were completed and 265 were not made. In Namibia, counts were made nearly annually between 1985 and 2019 at the four major colonies that constitute > 95% of the breeding population in that country: Mercury Island, Ichaboe Island, Halifax Island, and Possession Island (Kemper, 2015) (Figure S2, Appendix 1).

The methods used to count the numbers of occupied nest sites of African penguins have been outlined in detail elsewhere (Shelton et al., 1984; Crawford et al., 2011). Briefly, counts were undertaken by teams of people walking through a penguin colony and counting occupied nest sites. Larger colonies were broken down into predefined census areas, each of which was counted separately. Counts in South Africa were predominately made between February and September each year, those in Namibia mainly between September and February (Crawford et al., 2011; Crawford et al., 2013). At some small and difficult to access localities counts made outside the main breeding seasons were used if no other count was available for that year. Where more than one count was made at a locality in a year, the highest count was taken to represent the number of pairs breeding that year (Crawford et al., 2011). An occupied site was considered active if it contained fresh eggs or chicks, or was defended by a non-moulting adult penguin, and considered potential if it was not active but showed recent signs of use, e.g. the presence of substantial fresh guano or nesting material, the recent excavation of sand from a burrow nest, the presence of many penguin footprints in its vicinity, or a combination of these factors. Breeding by African penguins is not always synchronous (Crawford, Shannon, & Whittington, 1999), so potential nests were counted as they may be occupied by pairs that have recently finished breeding or that are about to breed (Crawford et al., 2011). Groups of unguarded chicks (crèches) were divided by two to estimate the number of nest sites they represented (mean clutch size is ~1.8 eggs (Crawford et al., 1999; Shannon & Crawford, 1999)), with remainders taken to represent an additional site, e.g. crèches of five and six chicks would both be taken to represent three nests (Shelton et al., 1984).

### 2.2 Generation length

The generation length (*G*) for African penguins was calculated as:

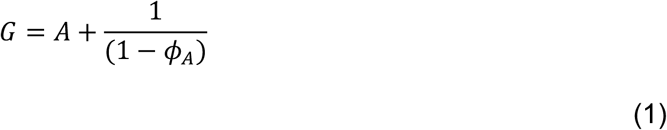

where *A* is age of first breeding and *ϕ_A_* is adult survival (BirdLife International, 2000). The IUCN Red List guidelines state “where generation length varies under threat … the more natural, i.e. pre-disturbance, generation length should be used” (IUCN Standards and Petitions Subcommittee, 2019). Accordingly we used *ϕ_A_* = 0.81 based on capture-mark-recapture studies at Robben and Dassen Island between 1989 and 1998 (Whittington, 2002) and between 1994/95 and 1998/99 (Sherley et al., 2014). African penguins can breed for the first time at between 4 and 6 years of age (Whittington, Klages, Crawford, Wolfaardt, & Kemper, 2005). Together these values yield generation length estimates of between 9.2 and 11.2 years. The previous assessment of African penguins used *G* = 10 years, and this value been supported by a recent meta-analysis of generation lengths in birds (Bird et al., 2020). Thus, we use *G* = 10 years here for consistency.

### 2.3 JARA state-space framework

To determine the trend and rate of change of the African penguin population we used JARA, a generalized Bayesian state-space tool for IUCN Red List assessments under Criterion A (Winker & Sherley, 2019) that has been applied recently to the Cape gannet *Morus capensis* (Sherley et al., 2019a) and several pelagic sharks (Sherley et al., 2019b). JARA assumes that the underlying trend in the population (*I_t_*) follows a conventional exponential growth model (Kéry & Schaub, 2012):

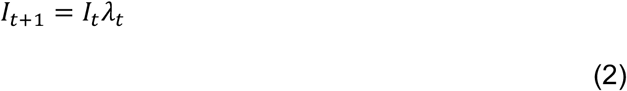

where *λ_t_* is the growth rate in year *t*. On the log scale, the process model was:

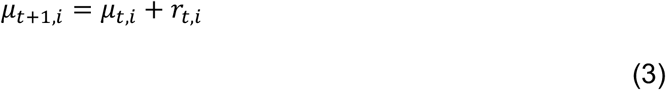

where *μ_t,i_* = log(*I_t,i_*) and *r_t,i_* = log(*λ_t,i_*), the year-to-year rate of change at breeding colony *i* that is assumed to vary around 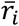 – the underlying mean rate of change for the colony – but with an estimable process variance 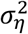 that is common to all colonies *r_t,i_*~Normal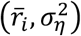. The corresponding observation equation is:

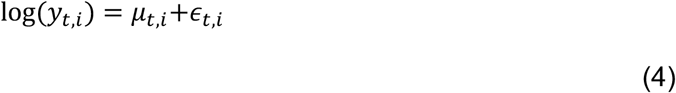

where *y_t,i_* is the number of pairs breeding in year *t* and *ϵ_t,i_* is the observation residual for year *t* at breeding colony *i*. The residual error is assumed to be normally distributed on the logscale *ε_t,i_*~Normal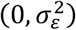 as a function of a common observation variance 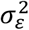, which is itself separated into two components: (1) a fixed input variance 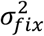 and (2) an estimable variance 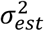. This fixed observation error accounts for additional sampling error associated with abundance indices and informs the process variance as a portion of total variance is assigned *a priori* to the observation variance (Winker, Carvalho, & Kapur, 2018; Winker & Sherley, 2019). Here, we set 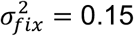 for all models. The estimated total population *Î_p,t_* for year *t* was then computed from the sum of all individual colony trajectory posteriors:

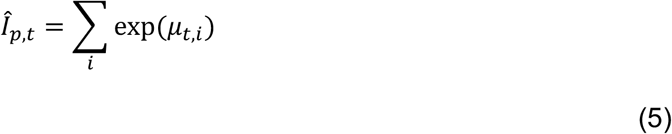

The change (%) in numbers at each colony was calculated from the posteriors of the estimated population trajectory (*Î_p,t_*) as the difference between a three-year average around the final observed data point *T*, and a three-year average around year *T* − (3G). The year *T* + 1 was projected to obtain a three-year average around *T* to reduce the influence of short-term fluctuations (Froese, Demirel, Coro, Kleisner, & Winker, 2017).

### 2.4 Regional variation in conservation status and decline rates

We first fit JARA simultaneously to the data from all 22 breeding colonies (18 in South Africa and 4 in Namibia) to determine the global trend, conservation status and rates of decline for the African penguin over the last three generation lengths (30 years). Thereafter, we subset the data and refit JARA to (1) the four Namibian colonies only to determine the trend, national status and rates of decline for Namibia; (2) the 18 South African colonies only, to give a perspective on the South African population. Then, to examine regional differences within South Africa, we further subset the data into (3) a West Coast region, in which we considered the seven South African colonies in the Western Cape that are north of Cape Town (Lambert’s Bay to Robben Island, Figure 1); (4) a South-West Coast region, which included the five Western Cape colonies south and east of Cape Town (Simonstown to Dyer Island, Figure 1); and (5) the six Eastern Cape colonies (Figure 1). To help model convergence, we used a value of 0.1 for the first year in time-series for colonies that were not yet occupied in 1979 (Robben Island, Stony Point and Simonstown) and for the last year at Lambert’s Bay as that colony went extinct in 2006 (Figure S1, Appendix 1).

### 2.5 Bayesian implementation

We implemented JARA in JAGS (v. 4.3.0) (Plummer, 2003) via the ‘jagsUI’ library (v. 1.5.1) (Kellner, 2017) for program R (v. 3.6.1) (R Core Team, 2018). The initials for the first modelled count *I_t=1,i_* were drawn in log-space from a ‘flat’ normal distribution with the mean equal to the log of the first observation *y_t=1,i_* and a standard deviation of 1000. We used vague normal priors of Normal(0,1000) for 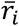 and inverse gamma priors for both the process and estimable observation variance 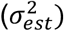 of *σ*^2^~1/gamma(0.001,0.001), which is approximately uniform on the log scale (Winker et al., 2018). We fit all models by running 4 Monte Carlo Markov chains (MCMC) for 400,000 iterations, with a burn-in of 200,000 and a thinning rate of 5. Convergence was diagnosed using the ‘coda’ package (Plummer, Best, Cowles, & Vines, 2006), adopting minimal thresholds of *p* = 0.05 for Heidelberger and Welch’s diagnostics (Heidelberger & Welch, 1992) and maximal thresholds of 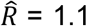 for Gelman–Rubin diagnostics (Gelman & Rubin, 1992). All models unambiguously converged. Unless otherwise specified, we report medians and 95% highest posterior density intervals (HPDI).

## 3. Results

### 3.1 Global population

Over the last 3 generations (30 years), the global African penguin population declined from ~53,300 pairs in 1989 to ~17,900 in 2019 (Figure 2a) at a median rate of change of −3.5% (HPDI: −6.5 to −0.5%) per annum (Figure 2b). This corresponds to a 64.1% (51.3–77.8%) decline, with 97.3% probability that the species meets the IUCN Red List classification of globally Endangered (EN) under criterion A2 (Figure 2c). The annual rate of change has remained around 4% since 1979 (All years = −3.5%, −6.5 to −0.5), but peaked at −5.6% (−9.2 to −2.1%) over the last 2G (Figure 2b).

**Figure 2.**
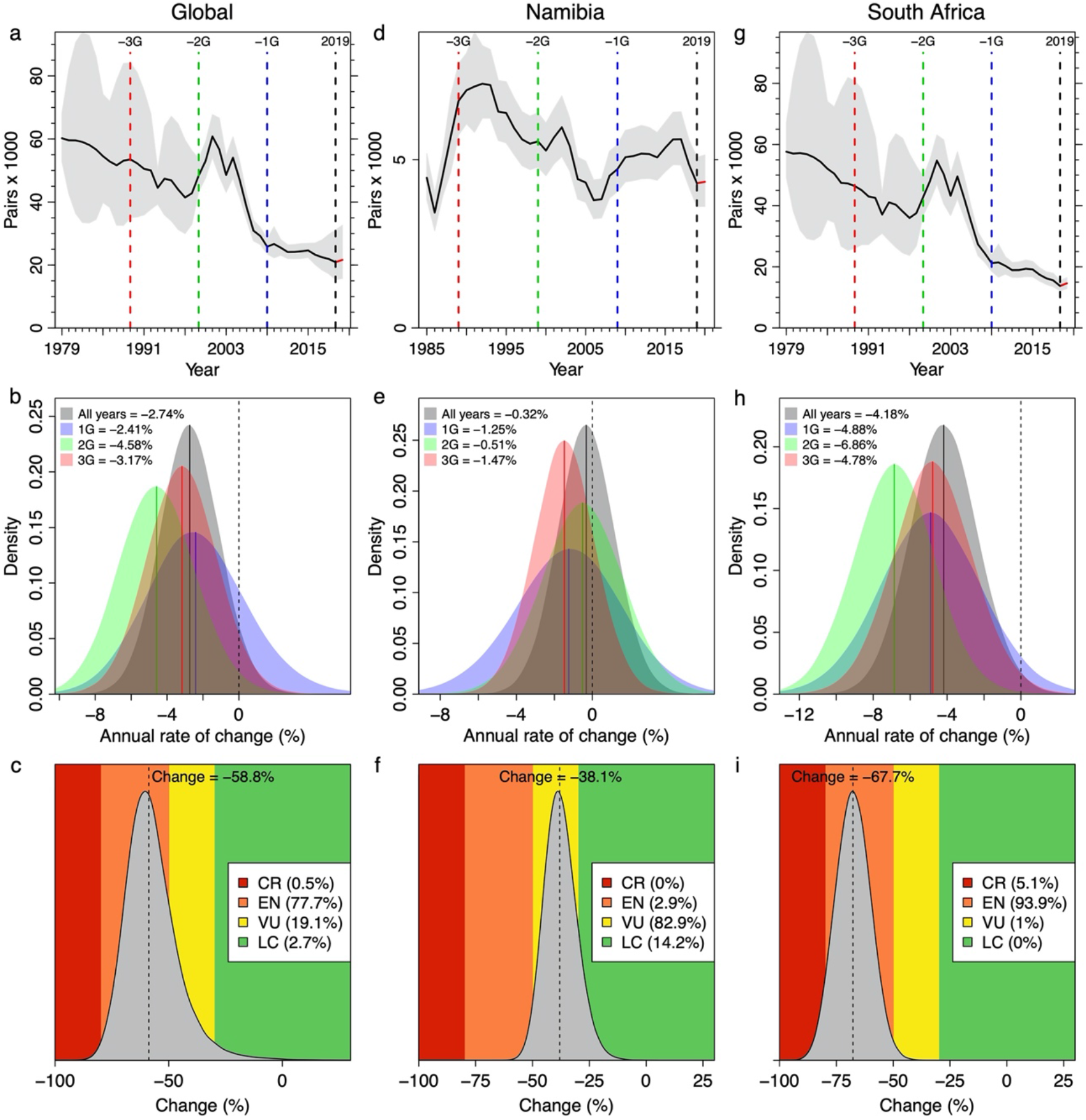
The decline in the global African penguin breeding population since 1979 (left panel), in the Namibian population since 1985 (centre panel), and in the South African population since 1979 (right panel). Top row (a, d, g): the JARA fitted median (black line) and 95% highest posterior density intervals (HPDI; grey polygon) for the population trend of African penguins based on nest counts from 22 colonies made between 1979 (1985 in Namibia) and 2019. The 10-year generation lengths before 2019 are denoted by a blue dashed line (−1G, 2009), a green dashed line (−2G, 1999) and a red dashed line (−3G, 1989). Middle row (b, e and h): the posterior medians (solid lines) and probability distributions (coloured polygons) for the annual rate of population change (%) calculated from all the data (All years, in black and grey), from the last one (1G; in blue), two (2G; in green), and three generations (3G; in red) shown relative to a stable population (% change = 0, black dashed line). Bottom row (c, f and I): the median change (%, dashed line) in the breeding population of penguins globally (c) in Namibia (f) and in South Africa (I) over three generations (3G) and corresponding posterior probability (grey polygon) for that change, overlaid on the IUCN thresholds for the Red List criterion A2 (LC—dark green, VU—yellow, EN—orange, CR—red).

### 3.2 Namibia – national status and trend

In Namibia, the African penguin population has fluctuated since 1985 (Figure 2d). Over the last 3G, however, the modelled population declined from ~6,700 pairs in 1989 to ~4,300 pairs in 2019 (Figure 2d). The median rate of change varied between −0.3 (−3.3 to +2.6) and −1.5 (−4.6 to +1.7)% (Figure 2e) as the population initially increased, then decreased through the 1990s and first half of the 2000s to a low of ~3,800 pairs in 2006, before recovering somewhat from 2008 (Figure 2f). Applying the IUCN Red List criterion A2 at a national level in Namibia would yield a classification of Vulnerable (VU) with a probability of 82.9% and a median decline over 3G of 38.1% (23.7–51.4%, Figure 3c).

**Figure 3.**
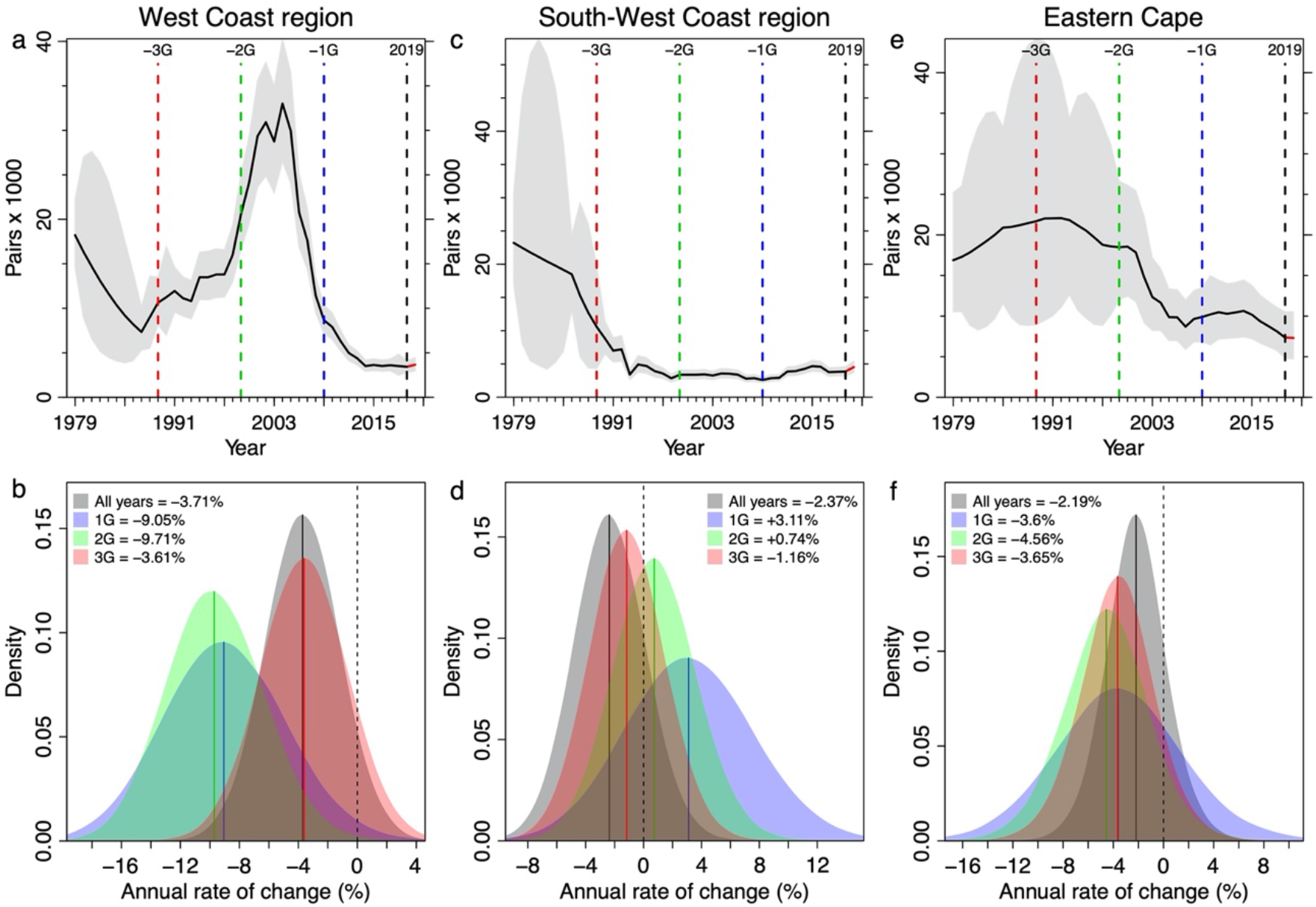
The change in the African penguin breeding population within the three regions of South Africa: the West Coast region (Western Cape colonies north of Cape Town; left, a and b), the South-West Coast region (Western Cape colonies south and east of Cape Town, middle, c and d) and the Eastern Cape (right, e and f). (a, c and e) The median (black line) and 95% HPDI (grey polygon) for the regional population trends of African penguins. The 10-year generation lengths before 2019 are denoted by a blue dashed line (−1G, 2009), a green dashed line (−2G, 1999) and a red dashed line (−3G, 1989). (b, d and f) The posterior medians (solid lines) and probability distributions (coloured polygons) for the annual rate of population change (%) calculated from all the data (1979 to 2019, All years, in black and grey), from the last one (1G; in blue), two (2G; in green), and three generations (3G; in red) shown relative to a stable population (% change = 0, black dashed line).

### 3.3 South Africa – national status and trend

Aside from a period of recovery during the late 1990s and early 2000s, the population in South Africa decreased fairly consistently since 1979 (Figure 2g), with an annual rate of change of −4.8% (−8.9 to −0.5%) over the last 3G (30 years, Figure 2h). Because of that period of recovery, the rate of change was fastest over the last 2G (20 years) at −6.9% (−11.1 to −2.7%), but the population continued to decline at −4.9% (−10.3 to +0.7%) per annum over the last 1G (10 years, Figure 2h). Applying the IUCN Red List criterion A2 at a national level in South Africa would yield a classification of EN with a probability of 93.9% and a median decline over 3G of 67.7% (54.5–82.3%, Figure 2i).

### 3.4 Regional trends within South Africa

Within South Africa, the bulk of the recovery seen in the national trend (Figure 2g) resulted from growth in the population in the West Coast region (Figure 3a) mainly Dassen Island and Robben Island (Figure S1, Appendix 1). Again, in part because of that period of growth and recovery, the rate of decline over the last 2G (20 years) has been substantial, at −9.7% (−16.0 to −3.3%, Figure 3b). However, unlike elsewhere, this rapid decline persisted in recent years; the rate of change at the colonies in the West Coast region over the last 10 years (1G) was −9.1% (−17.0 to −0.9%, Figure 3b). Overall, this regional population has declined by 68.7% (58.5–77.3%) at an annual rate of change of −3.6% (−9.3 to +2.1%) per annum over the last 30 years. Moreover, there was little uncertainty in this decline; if the IUCN Red List criterion A2 were applied at a regional level, this sub-population would qualify for an EN status with 99.7% probability (Figure S2, Appendix 1).

The trend at colonies in the South-West Coast region was initially dominated by a decline at Dyer Island, from ~23,000 pairs in 1979 to ~2,300 pairs in 1999 and ~1,050 pairs in 2019 (Figure S1, Appendix 1); thus the median rate of change since 1979 was −2.4% (−7.1 to +2.6%) overall and −1.2% (−6.1 to +4.0%) since 1989 (3G, Figure 3c). More recently, the decreases at Dyer Island were somewhat offset by the colonisation and growth (since the 1980s) of the land-based colonies at Simonstown and Stony Point to ~975 and ~1,750 pairs respectively (Figure S1, Appendix 1). As these two colonies have come to dominate the population numbers in this region, so the annual rate of change has shifted from negative to positive, ending at +3.1% (−5.3 to +11.8%) in the last 1G (Figure 3d). However, these increases did not offset the ~90% decline of the population at Dyer Island (Figure 3c).

In the Eastern Cape, the population has decreased fairly consistently since 1989 (Figure 3e) at rate of change ranging from −3.6 (−9.8 to +2.1) to −4.6 (−11.6 to +2.1)%, which has in general been slightly slower than the overall rate of change in South Africa (cf. Figure 3f with Figure 2h). Although this sub-population has declined by 66.4% (35.9–88.8%) over the last 3G (Figure S2, Appendix 1), it has come to represent a far greater proportion of the overall African penguin population in South Africa as a result of the substantial declines at Dyer Island and the colonies north of Cape Town (in particular Dassen Island). In 1979 the six Eastern Cape colonies contained 26 (19–34)% of the total African penguin population. In 2019, they contained 41 (30–53)%.

## 4. Discussion

African penguin numbers declined steadily over their last three generations, resulting in a loss of almost 65% since 1989, and reached an historical low of ~17,900 pairs in 2019. Our results strongly support its classification as globally Endangered on the IUCN Red List and indicate a clear cause for concern for this species. However, our use of JARA allowed us to decompose this decline in both space and time, and to demonstrate robustly that African penguins have not declined uniformly across their range. This variability has arisen for several reasons, including differences in the nature and severity of threats and local population dynamics. It follows, then, that there are different conservation management priorities for each subpopulation.

The Namibian population has declined at a slower rate than the other regional populations over the last three generations, with the rate of decline sufficient to warrant a Red List classification of Vulnerable under criterion A (Figure 2f). However, the Namibian penguin population had already declined by ~70% prior to the start of our dataset in 1986, coincident with the collapse of the Namibian sardine stocks in the 1970s (Crawford, 2007). The population also underwent a worrying decline to 3,600 pairs in 2007 before recovering slightly to 4,300 pairs by 2019. The penguin population in Namibia is likely now constrained at a low number by a scarcity of small pelagic fish (Watermeyer, Shannon, Roux, & Griffiths, 2008; Roux et al., 2013) and the birds’ reliance on lower energy prey (Ludynia, Roux, Jones, Kemper, & Underhill, 2010). Monitoring of breeding colonies in Namibia is an ongoing priority, with an annual census of breeding pairs the minimum requirement to track trends in this population. A recent outbreak of avian influenza in some colonies in Namibia has shown the vulnerability of this population to stochastic events (Molini et al., 2020), the effects of which are exacerbated at low population levels (Lande, 1993).

The South African population recently declined at a much faster rate than the one in Namibia, resulting in a national and global classification of Endangered. Despite a small population recovery in the late 1990s and first half of the 2000s, driven mostly by increases in the West Coast region, there was a subsequent crash from the mid-2000s onwards to an historical low in South Africa of ~13,700 pairs in 2019. The short-lived population recovery and subsequent crash were associated with a concomitant boom and then decline in sardine and anchovy biomass (Crawford et al., 2011). This decline also coincided with an eastward displacement of a number of marine resources in South Africa (Blamey et al., 2015), including spawning adults of sardine and anchovy (Roy, van der Lingen, Coetzee, & Lutjeharms, 2007; Coetzee, van der Lingen, Hutchings, & Fairweather, 2008). These environmental changes combined with fishing pressure (Coetzee et al., 2008; Mhlongo, Yemane, Hendricks, & van der Lingen, 2015) to lower the availability of prey for seabirds breeding to the north of Cape Town (Crawford, Sydeman, Thompson, Sherley, & Makhado, 2019). This loss of their prey base underpinned the dramatic and unsustainable decline at almost 10% per year over the last 20 years at the West Coast colonies (Figure 3b). In contrast, penguin numbers in the South-West Coast region have remained relatively stable at low levels over the last three generations, principally supported by growth of the mainland colonies at Simonstown and Stony Point, which has somewhat offset the recent portion of the long-term decline at Dyer Island (Figure S1). Meanwhile, the Eastern Cape region has experienced periods of relative stability followed by declines in the early 2000s and the late 2010s. Because the Eastern Cape population has declined at a slower rate than elsewhere in South Africa, the area has become increasingly important in terms of its relative contribution to the global population. At the same time, Algoa

Bay has been identified as a marine transport hub, with permitted ship to ship bunkering taking place, and potentially as an Aquaculture Development Zone (Massie et al., 2019), increasing the risks of oil spills and other human impacts on the ecosystem (Pichegru, Nyengera, McInnes, & Pistorius, 2017; Ryan, Ludynia, & Pichegru, 2019). Since bunkering was permitted in Algoa Bay, for example, two bunkering related oil spills have taken place, in 2016 and 2019, oiling 220 African penguins (Ryan et al., 2019).

A lack of suitable prey, predominantly small pelagic fish, is believed to be the main driver for declines in African penguin numbers in South Africa over the last three generations (Crawford, Sabarros, Fairweather, Underhill, & Wolfaardt, 2008; Crawford et al., 2011; Sherley et al., 2017; Crawford et al., 2019), with sporadic oiling events, habitat destruction, disturbance, and predation also contributing to the losses (Crawford et al., 2000; Pichegru, 2012; Makhado, Crawford, Waller, & Underhill, 2013; Weller et al., 2014). In 2013, the South African government put in place a Biodiversity Management Plan (BMP) for the African penguin (DEA, 2013). This plan aimed to halt the decline of the species and thereafter achieve the down listing of the species’ conservation status. Although the plan did not achieve its aim, it provided a coordinated approach to penguin conservation and several key conservation interventions were initiated, or given greater credence, through this plan. One conservation intervention given increased importance in the BMP is the identification and protection of important foraging areas. Work along these lines has focused predominately on a 12-year experiment, started in 2008, to investigate the effects of fishing closures around penguin breeding colonies. The experiment has shown some benefits to breeding penguins through a decrease in foraging effort and an increase in chick growth and condition when fishing was prohibited (Pichegru et al., 2012; Sherley et al., 2015, 2018) (although this has been contested, Butterworth, Plagányi, Robinson, Moosa, & Moor, 2015; Robinson, Butterworth, & Plagányi, 2015; Weller et al., 2016). The recent stability of breeding numbers at Simonstown (small pelagic fishing in False Bay has been prohibited since 1982, Penney, 1991) and Stony Point (which is surrounded by a small marine protected area) during a period when the populations at all the other South African colonies have declined, also provides circumstantial evidence in support of protecting the key foraging areas used by breeders.

The initial identification of areas used by penguins during other parts of their life cycle such as pre- and post-moult and during the first few years after fledging has also begun (Roberts, 2016; Sherley et al., 2017), but further work is required to determine the most appropriate mechanism to protect penguins during these vulnerable periods (Sherley et al., 2017). Additional spatial management of sardine and anchovy fishing effort, currently concentrated on the West Coast, will assist with addressing the mismatch between fish distribution and fishing effort (Coetzee et al., 2008; Grémillet et al., 2008). The hand-rearing and release of chicks (Sherley et al., 2014), and the creation of new breeding colonies have also been suggested as ways to mitigate the mismatch between penguin breeding colonies and fish distribution (DEA, 2013) and a pilot site to establish a colony is currently underway on the southern coast of South Africa. A revised BMP is being prepared with fewer, more threat-focused actions, and will be implemented from 2020.

Our results highlight the dynamic nature of the decline of the African penguin population and have clarified the long-term regional population trajectories. We identified an unsustainable decline of almost 10% per year at colonies to the north of Cape Town, the former geographic core of the species’ breeding range. Our results denote a shift to a condition where colonies at the geographic edge of the species’ range in the Eastern Cape form the stronghold of the African penguin population, a situation analogous to the poleward shifts seen in marine taxa elsewhere (Hastings et al., 2020). Accordingly, the Eastern Cape colonies should be viewed as a priority for conservation interventions, as should actions that could contribute to retaining viable breeding populations at the formerly large colonies in the West Coast region (Sherley et al., 2018). Finally, the robust JARA-based Red List Assessments we used provide transparent estimates of uncertainty, accommodate non-linearity in population trajectories, and account for observation heterogeneities (Sherley et al., 2019b). Thus, our approach has wide potential applicability to other studies of wildlife populations threatened with extinction.

## Acknowledgements

We thank CapeNature, SANParks, the South African Navy, Raggy Charters, Robben Island Museum and our institutions for logistical support, and the many people who have helped with counting penguins since 1979. R.B.S. was supported by the Pew Fellows Program in Marine Conservation at The Pew Charitable Trusts. The views expressed are those of the authors and do not necessarily reflect the views of The Pew Charitable Trusts. This paper is an output of the Benguela Current Commission’s BECUMATOP programme.

## Author contributions

R.B.S. and H.W. conceived the study and designed the analysis; R.B.S., R.J.M.C., B.M.D., D.G., J.K., A.B.M., L.P., D.T., L.U., J.V. and L.J.W. contributed to data collection; R.J.M.C., J.K., A.B.M. and D.T. collated the data; R.B.S. made figure 1, and analysed the data and made the remaining figures using JARA code written by H.W. and R.B.S.; R.B.S., A.D.d.B. and C.H. wrote the first draft of the manuscript; All authors contributed to revisions and gave final approval for publication.

## Data availability statement

The data underlying this article will be made available on Dryad upon acceptance for publication.

## Competing interests

The authors declare no competing interests.

## Appendix 1

**Figure S1.**
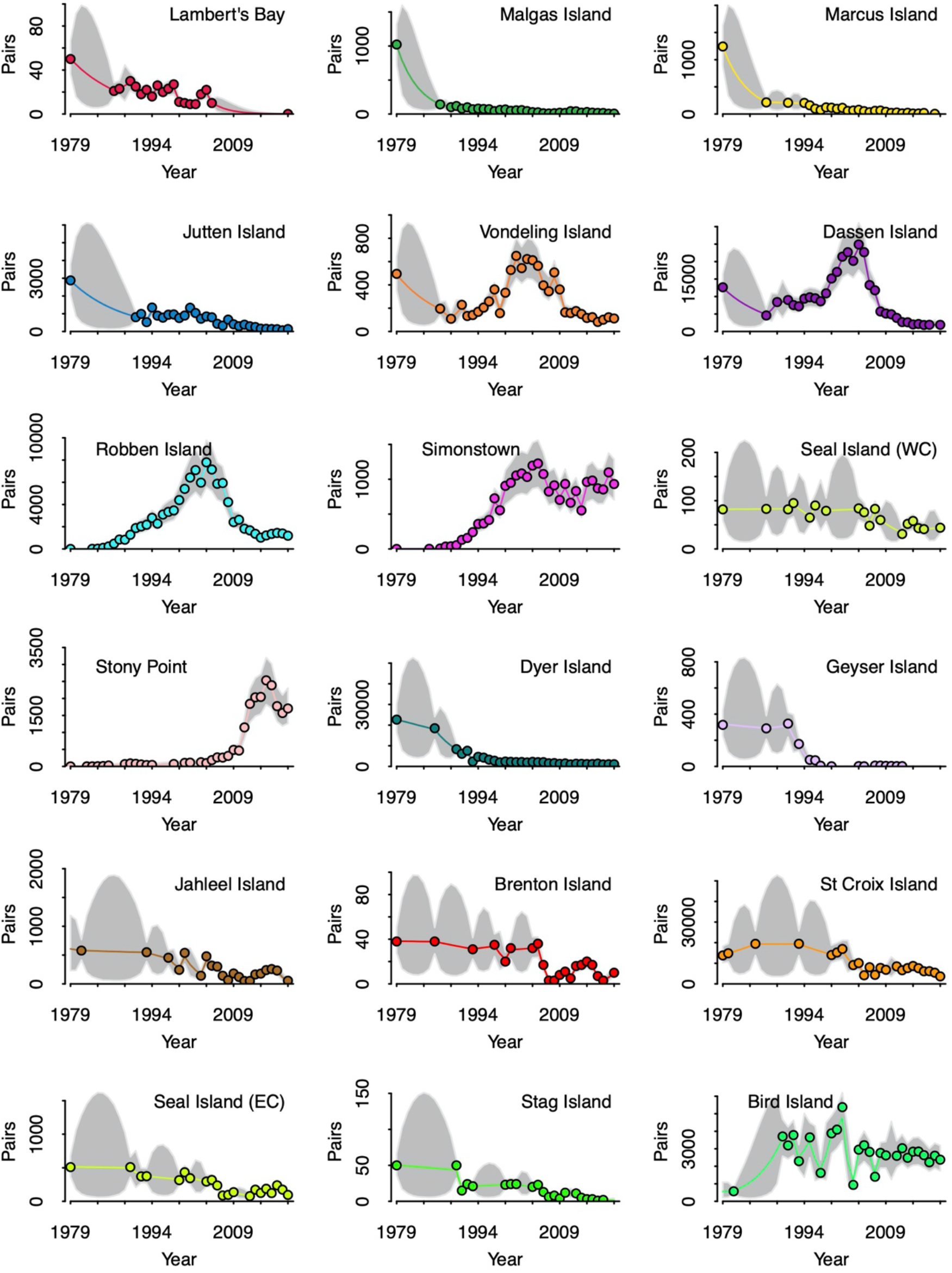
Bayesian state-space model fits (lines) from Just Another Red List Assessment (JARA) to population counts (points) made between 1979 and 2019 at 18 of the 19 known colonies in South Africa at which African penguins have bred for more than 5 years during that time frame.

**Figure S2.**
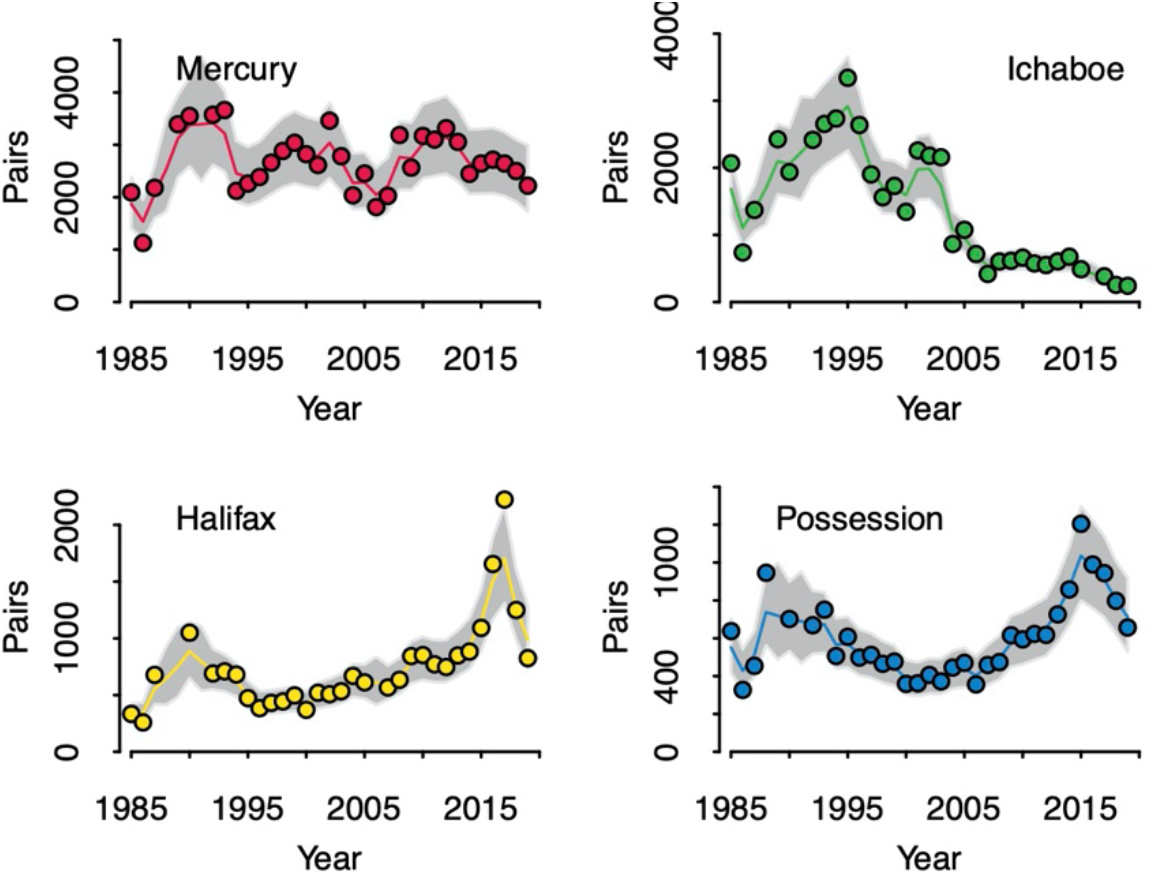
Bayesian state-space model fits (lines) from Just Another Red List Assessment (JARA) to population counts (points) made between 1985 and 2019 at the four breeding colonies in Namibia that contained ~95% of the Namibian African penguin population since 1985 (Kemper, 2015).

**Figure S3.**
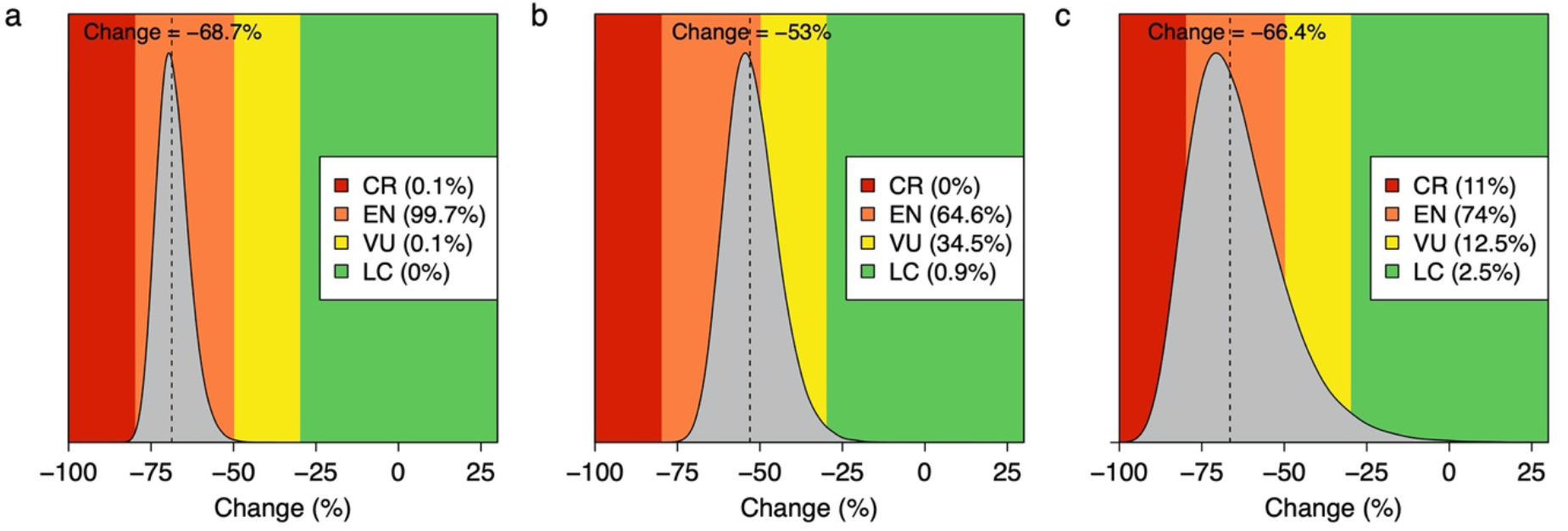
The median change (%, dashed line) in the breeding population of penguins in (a) the West Coast region of South Africa, (b) the South-West Coast region of South Africa, and (c) the Eastern Cape province of South Africa over three generations (3G) and corresponding posterior probability (grey polygon) for that change, overlaid on the IUCN Red List category thresholds for the Red List criteria A2 (LC—dark green, VU—yellow, EN—orange, CR—red).

## Notes

### Competing Interest Statement

The authors have declared no competing interest.

### Summary of Updates

Reanalysis after receiving updated data from Namibia. Wholesale revision of text based on new results. Final version before submission to a peer-reviewed journal.

